# A novel CT_max_ assay reveals divergent thermal acclimation capacity across three ecologically distinct sea urchins

**DOI:** 10.64898/2026.08.19.745849

**Authors:** Daniel E. Sadler, Andrew R. McCracken, Caroline Deir, Cormac Bassett, Tran B. Vu, Joaquin C.B. Nunez, Melissa H. Pespeni

**Affiliations:** Department of Biology, University of Vermont, Vermont, U.S.A

**Keywords:** Thermal tolerance, CTmax, Sea Urchins, Climate change, Echinoderms

## Abstract

Global change is driving rapid ocean warming, exposing organisms to both chronic temperature increases and acute marine heatwaves. Understanding how species cope with thermal stress is critical for predicting ecosystem resilience. Echinoderms are globally distributed and often function as foundational species, yet comparative assessments of upper thermal tolerance among species occupying contrasting thermal environments remain limited. Here, we address this gap by comparing upper thermal tolerance across three sea urchins with distinct biogeographic distributions: the latitudinally broad purple sea urchin (*Strongylocentrotus purpuratus*), the circumpolar green sea urchin (*S. droebachiensis*), and the tropical variegated sea urchin (*Lytechinus variegatus*). We quantified thermal limits after two acclimation treatments: ambient temperatures approximating native habitat conditions for each species and an elevated temperature (+6 °C). We developed a novel assay to measure critical thermal maximum (CT_max_), comparing variability and inconsistencies associated among multiple assays. Upper thermal tolerance increased with acclimation to elevated temperatures in all three species, but acclimatory capacity differed markedly, with *S. droebachiensis* showing the strongest response and *S. purpuratus* the weakest. Conversely, *S. purpuratus* had the highest thermal safety margin and thus the lowest proximity to its thermal ceiling. Our adhesion based CT_max_ method was more reproducible and the most precise compared to other metrics tested, providing an improved framework for quantifying physiological thermal limits of sea urchins. Together, these findings reveal substantial but unevenly distributed thermal resilience in ecologically diverse sea urchins, advancing our understanding of how foundational marine species may respond to future global change.

**Summary Statement:** Using a novel CT_max_ assay, we find thermal acclimation increases upper heat tolerance, but unevenly, across three sea urchin species from distinct ecological niches.

## 1. Introduction

### Global change and its influence on marine organisms

Global change is driving rising sea surface temperatures worldwide through both long-term ocean warming and an increasing frequency of short-term extreme events, such as marine heatwaves, which are becoming more frequent and intense (Oliver et al., 2018; Sen Gupta et al., 2020). Marine ectotherms are particularly vulnerable to these shifts because their physiological performance is closely tied to environmental temperature (Pörtner and Farrell, 2008), and rising temperatures accelerate biochemical processes and increase metabolic rates, with downstream effects on growth, development, reproduction, and survival (Huey et al., 2001; Lang et al., 2023; Zuo et al., 2012). Whether a species can buffer these effects depends in part on its physiological capacity to acclimate to short-term warming, but this capacity varies widely across taxa and is shaped by a species’ thermal history (Morley et al., 2019). Because acclimation capacity determines how well populations withstand acute heat events layered on top of long-term warming, differences among species could have lasting consequences for population fitness and ecosystem health (Stillman 2003; Somero 2010; Wernberg et al., 2025), particularly for foundational species, whose decline can reshape entire communities.

### Thermal adaptation across geographic ranges

As global change alters temperature regimes, species will be affected differently depending on their current biogeographic range and associated thermal capacities. Thermal tolerance to rising temperatures is closely associated with the magnitude of thermal variation experienced by populations in their native environments (Hughes et al., 2019; Khaliq et al., 2014; Sunday et al., 2010); and as such is often generally associated with latitude following the climate variability hypothesis (Janzen, 1967; Stevens, 1989). Therefore, stenothermal species such as tropical and subarctic species will likely be particularly vulnerable to future climate fluctuations, with even moderate changes in temperature acclimation having severe fitness consequences (Richard et al., 2012; Sørensen et al., 2024; Tewksbury et al., 2008). Beyond current thermal tolerance, it is important to know how close populations and species are to their thermal limits, and whether they can acclimate to future climatic change (Stillman, 2003). Populations near their thermal limit have a reduced capacity to acclimate, and is evident even within the same species, for example, porcelain crabs showed reduced acclimation to thermal stress in warm-adapted populations (Stillman, 2003) compared to cool-adapted populations. Local thermal variability may therefore select for broad tolerance and high plasticity in variable environments or narrow tolerance and high performance in stable ones (Somero, 2010; Sunday et al., 2012). As contemporary global change is occurring faster than adaptation, population resilience will strongly rely on acclimation capacity (Stillman 2003; Somero 2010). Understanding how thermal tolerance and acclimation capacity differ across species and populations from ecologically diverse habitats and evolutionary histories is important for predicting vulnerability to global change.

### Importance of sea urchins as a model/ foundation species

Echinoderms, and sea urchins specifically, are a key foundational species in habitats ranging from the high arctic to the tropics and can be considered sentinels of ecosystem health (Byrne and O’Hara, 2017; Pinsino and Matranga, 2015). For example, sea urchins in tropical coral reef systems graze on macroalgae, maintaining a healthy reef system (Fong et al., 2024; Williams, 2022). However, urchins can also overgraze algae, such as in kelp forest systems leading to urchin barrens (Filbee-Dexter and Scheibling, 2014), which have large-scale economic and ecological consequences (Eger et al., 2023, 2024). In addition, many urchin species around the world are harvested for their roe, supporting economically important fisheries (Andrew et al., 2002; Sun and Chiang, 2015). Previous studies in the purple sea urchin, *Strongylocentrotus purpuratus*, have shown that sea urchins have both genetic and physiological capacity to respond to environmental stress (Pespeni et al., 2013, 2012, 2010; Pespeni and Palumbi, 2013; Petak et al., 2023), potentially linked to their high standing genetic diversity (Pespeni et al., 2012, 2010; Petak et al., 2023). This level of resilience is likely due to the high levels of gene flow and the high degree environmental variation experienced across both space and time for this temperate species (Flowers et al., 2002; Pespeni et al., 2010; Pespeni and Palumbi 2013). Species that experience less environmental variation, such as those in polar or tropical environments, may be less resilient (Lang et al., 2023; Morley et al., 2019; Sunday et al., 2010). Given their ecological and economic importance and broad distribution, understanding sea urchin thermal tolerance is critical for predicting ecological dynamics in future global change conditions.

### CT_max_ as a metric for future resilience to change

Critical thermal maximum (CT_max_) is a metric often used in physiological studies of thermal tolerance; it is the temperature at which an organism loses motor function during rapid warming (Cowles and Bogert, 1944). CT_max_ is typically used for ectotherms including fish (Desforges et al., 2023), but is less prominent in aquatic invertebrates (Bayat et al., 2025; Cereja, 2020), where it is more difficult to quantify loss of motor response or failure of predator escape. CT_max_ is also commonly used to measure whole organism acclimation, (Comte and Olden, 2017; Ruthsatz et al., 2024) and has been correlated with long-term warming (Åsheim et al., 2020) and mortality of species under prolonged thermal stress (Cicchino et al., 2023). Additionally, the rapid ramping temperatures of a CTmax assay are similar to acute warming events, for example, mid-day low tides for intertidal species during a hot summer day. In general species from tropical, warmer habitats have greater CT_max_ than temperate and polar species (Sunday et al., 2012) however, ability to shift CT_max_ under acclimation to higher temperatures is often lower for tropical species, as they are already close to their thermal ceiling. Assessing both acute thermal stress and general patterns of thermal adaptation using CT_max_ is useful for comparing vulnerability across taxa.

### CT_max_ in echinoderms requires a consistent, repeatable protocol

A key consideration when measuring CT_max_ is the variability in methodology across studies (Raby et al., 2025). However, marine invertebrates, in particular echinoderms, have relatively few studies that assess CT_max_. This may in part be due to the challenge of determining the end point, i.e., the maximum temperature, for urchins. For example, thermal assays in fish or flies often have a clear end point, loss of equilibrium (knockdown) or cessation of movement despite gentle prodding. In previous studies of sea urchins, several methods have been used to assess whole organism performance: righting response, i.e., time for urchin to flip itself from an upside down position to normal orientation at a given temperature (Percy 1973), righting time at the temperature 1°C below the empirically determined mortality temperature (Collin et al., 2018), and CT_max_, with progressive temperature increases, the temperature at which tube feet retract and spines compress (DÍaz et al., 2017; Hernández et al., 2004) or spines and tube feet become immobile (Lemus-Granados et al., 2025). In our preliminary studies, however, we found inconsistencies within these measures between individuals and species and between replicate assays, with some measures potentially being subjective and prone to human error or representative of behavioural traits rather than a physiological limit. As such there is a need to develop a methodology that is consistent and repeatable across sea urchin taxa. Given the common definition of CT_max_, the temperature at which an organism loses motor function and is unable to escape predation (Cowles and Bogert, 1944; Raby et al., 2025), we propose adhesion to substrate, essential for avoiding predation and staying fixed to the substrate despite wave action, as a similar proxy for urchins and relevant metric for their CT_max_.

To understand the influence of thermal acclimation on upper thermal tolerance (measured as CT_max_), we used three species of sea urchin from a diverse range of habitats: the subarctic green sea urchin (*Strongylocentrotus droebachiensis*, O.F. Müller, 1776), the latitudinally broad purple sea urchin (*Strongylocentrotus purpuratus,* Stimpson, 1857), and the tropical variegated urchin (*Lytechinus variegatus*, Lamarck, 1816). Urchins were acclimated to ambient (control) and elevated (+6°C) temperatures, a temperature indicative of seasonal highs across their species ranges and representative of long-term extreme heatwave events (Sen Gupta et al., 2020). In assessing CT_max,_ tube feet retraction, spine compression, and righting response among groups, we addressed the following questions (1) Do ecologically distinct sea urchins differ in acclimation capacity? (2) How does acclimation temperature affect thermal tolerance? And (3) Can we develop a standardised, reproducible thermal assay?

## 2. Methods

### 2.1. Experimental Design

Green sea urchins (*S. droebachiensis*; n = 40), variegated urchins (*L. variegatus*; n = 40) and purple sea urchins (*S. purpuratus*; n = 20) were collected and used for the study. The green urchins were obtained from the University of Maine Center for Cooperative Aquaculture Research from lab bred lines collected from the gulf of Maine and maintained at 8°C. The purple sea urchins were collected at Bodega Marine Station, by UC Davis (38.32° N, 123.07° W) and maintained at 12°C. The variegated sea urchins were obtained from the Florida keys (Carolina Biological Supply) and maintained at 21°C. All urchins were maintained in artificial sea water in separate 190 L stock tanks. After two weeks of acclimation to stock temperatures, urchins were moved to adjacent 190 L tanks randomly assigned to four tanks of two temperature treatments (ten urchins per tank [five for *S. purpuratus*]): control (ambient stock temperature) and thermal stress (+6°C from ambient). The thermal stress treatment was achieved by ramping the temperature up 1°C a day. Ambient temperature of each species is representative of their native environment, whilst a +6° is representative of acute thermal stress and the increase is comparable to the upper end of predicted heatwave scenarios (Sen Gupta et al., 2020). All urchins were acclimated for four weeks under these temperature conditions. Temperature and salinity were monitored daily, whilst nitrate levels were monitored weekly to ensure environmental stability. Urchins were fed with Kombu dried seaweed (Emerald Cove) weekly before the experiment began, then were starved for the duration of the thermal acclimation to standardise physiological activity and maintain water quality throughout the experiment.

Wet weight (g), and length (mm) was measured once a week per individual. Wet weight was taken by placing an urchin in a weighing boat on a scale (Mettler Toledo, U.S.A), whilst length was taken by using callipers on the test. However, due to the inaccuracy of length measurements due to spine movement, we decided to use only wet weight to allow correction for size-related performance effects. Additionally, colour morph of the variegated urchin was recorded to test for any potential differences in thermal tolerance based on colour (Wise et al., 2024).

### 2.2. Righting Response

Righting response was measured weekly for green and variegated urchins. For righting response, each urchin was placed into a large 2 L glass beaker filled with water from its origin tank. The individuals were timed in seconds from time of placement at the bottom of the beaker to the time it flipped itself to halfway point (90°). Water was replaced after each trial and urchins placed in a recovery tank before being placed back into the origin tank to prevent any influence on other urchins in the tank. Righting response was converted to an activity coefficient (AC) using the formula (AC = 1000/righting time− 600/1000), with a lower AC indicating a longer time to right and poorer performance and higher AC indicating a shorter time to right, and failure to right within the 600s becomes zero (Barker and Russell, 2008).

### 2.3. CT_max_ Trial

To prepare for the CT_max_ trials, each urchin in both tanks (n = 10 per tank) was assigned a unique coloured zip tie for individual identification based on wet weight and length. The zip ties secured the urchins inside 5 × 5 in. flow-through cages within each tank (**Figure 1**). Four of the sides were made of perspex that could be adhered to by the sea urchin, whilst two of the sides were made from custom cut aquaria grating with 10 mm x 10 mm grids (Alegi, China) to allow for water flow. Full list of components for the CT_max_ system in **Table S1**.

**Figure 1:**
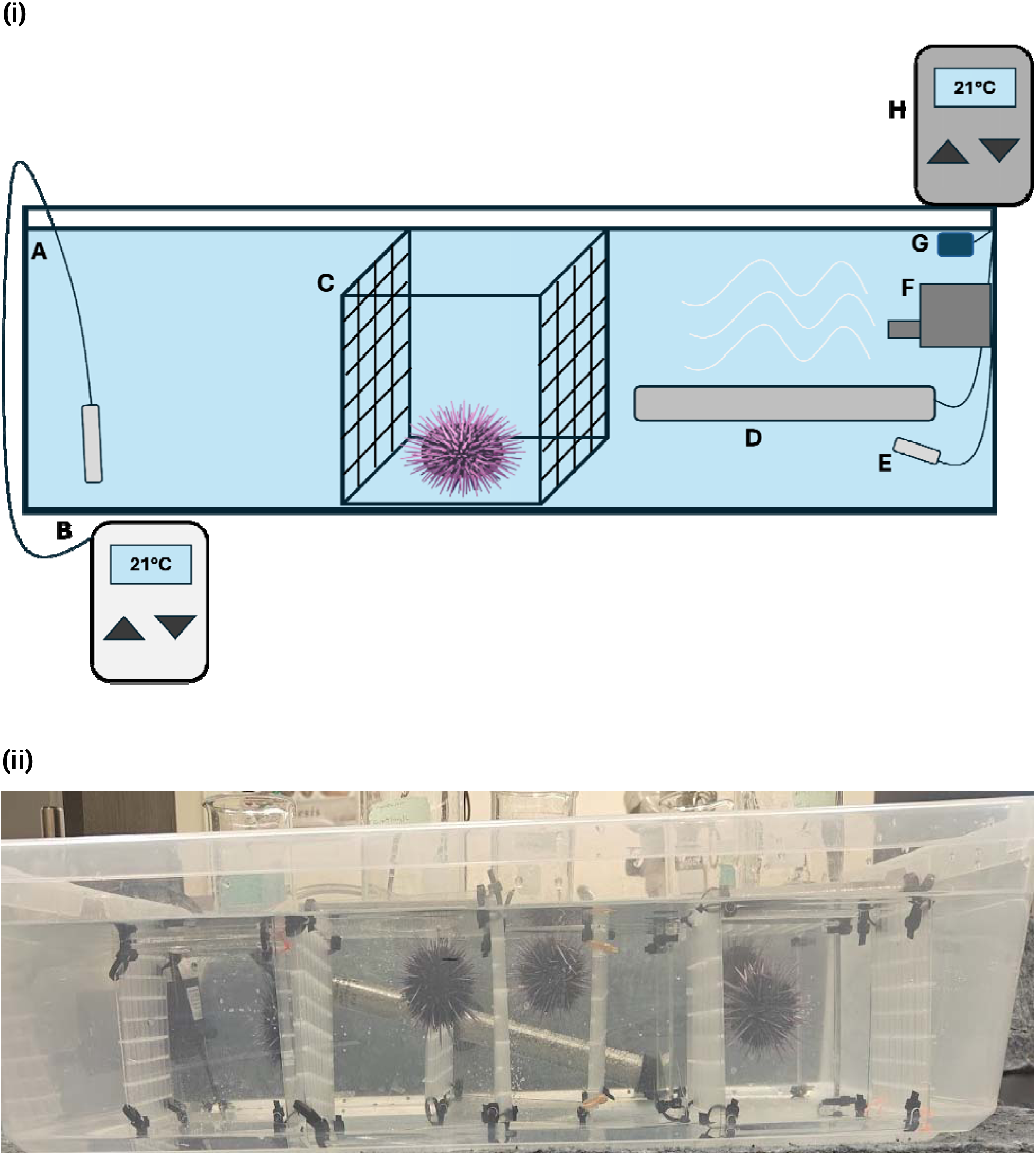
Experimental setup for the CT_max_ experiment. (i) Mock example set up for one cage: (a) Plastic tub containing 100L saltwater, (b) Testo temperature probe, (c) plastic cage with flowthrough holes, (d) titanium heater, (e) temperature probe, (f) water pump, (g) airstone, (h) inkbird temperature controller. (ii) Side-view photo of the assay system set up for all five urchins numbered accordingly.

CT_max_ was assessed once in week five for *S. purpuratus* and twice for *S. droebachiensis* and *L. variegatus*, once in week five and once in week six. Each trial was conducted over two days, with one ambient tank and one elevated temperature tank tested per day, within each temperature treatment, temperatures were ramped up one day apart between the replicate tanks to stagger the two tanks, as it was not possible to complete all CT_max_ assays across 40 urchins in one day. Instead, we divided the CT_max_ across two days to allow for the same time between trials per tank. We randomly assigned whether a tank replicate was measured either in the morning or afternoon and switched for the subsequent trial to account for any effects of photoperiod. For each run, urchins were identified by their colour tag, removed from their cages, weighed, and then placed individually into flow-through chambers inside a 30 L open-top plastic container which was wrapped in opaque insulative material (**Figure 1**). Five urchins were tested at a time to ensure accurate observation of each. Two observers were always present and watching from above for the duration of the assay. Individuals acclimated at their respective treatment temperature for 15 minutes in the container before heating began. Water temperature increased at a rate of 0.3°C per minute using a titanium submersible heater connected to a temperature controller (Inkbird, Hong Kong). A water pump and air stone were used to ensure even heat and oxygen distribution. Temperature was additionally measured using a precision temperature probe (Testo, USA). Water was changed after every trial and urchins placed individually in 3L recovery beakers until all urchins of the origin tank were assayed, then returned to their original experimental tanks.

During heating, three phenotypic indicators of stress were recorded. The first was tube feet retraction, which is the temperature at which the urchin’s tube feet were no longer visible, retracting into the test. The second indicator was spine decompression, which was recorded when at least half of the spines folded over, making the urchin’s normally round shape look more pentagonal. The final indicator was the loss of adhesion (CT_max_), measured as the temperature at which the urchin detached from the cage and was unable to reattach. The cage was rotated whenever the urchin rested at the bottom or reattached itself after falling to confirm that adhesion loss was final. The number of rotations per individual was recorded to correct for any confounding effect of handling stress. Immediately after final loss of adhesion, each urchin was removed from the container and returned to its home tank. To track individuals, the colour identifiers were used in weeks five and six for both CT_max_ trials and the corresponding righting-response measurements. We maintained the urchins for a week past the trial to identify any mortalities, with one urchin (green sea urchin) dying 24 hours after the trial after achieving an unusually high CT_max_ (30°C at which point we halted the assays). In addition to CT_max_, we calculated thermal safety margin (TSM [mean CT_max_ - acclimation temperature]) and acclimation response ratio (ARR [change in thermal limit/change in acclimation temperature]; Claussen, 1977).

### 2.4. Statistical Analysis

All analyses were performed in R version 4.5.1 (R Core Team, 2025). To assess the effects of temperature on loss of adhesion, spine decompression, and tube feet retraction (proxies of CT_max_), we used a generalised linear model (GLM; CT_max_ proxy∼Treatment) with the purple sea urchin, whilst for the green and variegated urchin, as we also tested repeatability, we included urchin ID into the model as random factor, running LMMs as CT_max_ proxy∼Treatment+(1|Urchin ID). We made sure the model fit assumptions of normality and heterogeneity using qqnorm plots and a Shapiro-Wilks test (Shapiro and Wilks, 1965).

We also assessed if weight was correlated with any of the measured traits using Pearson’s correlation test, but did not have a significant correlation with adhesion, tube feet, or spine compression. Additionally, photoperiod (morning or afternoon assay) and number of cage rotations had no significant effect on CT_max_ so we excluded those variables from the model.

We additionally tested the effects of temperature and week on activity coefficient (AC). Due to non-normality of the righting response data, we used a generalized mixed effect model (GLMM) using template builder model with the package *glmmTMB v.1.1.14* with tweedie family and log link. Performing likelihood ratio tests, and comparing AIC, the final model we used was AC ∼ Treatment + Week + (1 | Urchin ID), family = tweedie(link = “log”) for both *L. variegatus* and *S. droebachiensis*.

Coefficients of variance values were calculated using the *cv* function in *EnvStats* v.3.1.0. We additionally tested repeatability across trials using the *rpt* function in *rptR v.* 0.9.23, as well as assessing correlations between trials with individual level repeats using Pearson’s correlation test.

## 3. Results

### 3.1. Loss of adhesion as a measure of CT_max_

Elevated acclimation temperature increased thermal tolerance (CT_max_), measured as loss of adhesion, in *S. purpuratus* (*F*_1,18_ = 6.32, *P-value* <0.05; **Figure 2; Table S2**), *S.droebachiensis* (*F*_1,37.6_ = 74.84, *P-value* <0.001; **Figure 2; Table S2**) and *L. variegatus* (*F*_1,39.5_ = 56.18, *P-value* < 0.001; **Figure 2; Table S2**). *S. purpuratus* had a marginal 3.57% (+1°C) increase in CT_max_ when acclimated to +6°C, whilst *S.droebachiensis* and *L. variegatus* had stronger responses with a 16.9% (+4.4°C) and 7.3% (+2.4°C) increase in CT_max_, respectively. We also found an increase in CT_max_ when using both spine compression and tube feet as proxies for all species (**Table S2**). To assess how close the organisms were to their thermal limit, we calculated thermal safety margin (TSM = CT_max_ – acclimation temperature). *S. purpuratus* had the highest TSM under ambient conditions (16°C), followed by *S. droebachiensis* (13.7°C), and the variegated urchin (11.9°C) which tracks with expectations of tropical species being closer to their thermal ceiling. However, when acclimated to an elevated temperature the TSM of all species decreased, and *S. droebachiensis* had the highest TSM at 12.1°C followed by *S. purpuratus* (11°C) and *L. variegatus* (8.3°C) urchins. Additionally, the acclimation response ratio (ARR; Claussen, 1977), the change in an organism’s thermal tolerance over the difference in acclimation temperatures, was highest for *S.droebachiensis* at 0.733 followed by *L. variegatus* (0.4) and *S. purpuratus* (0.166) (**Figure 3**).

**Figure 2:**
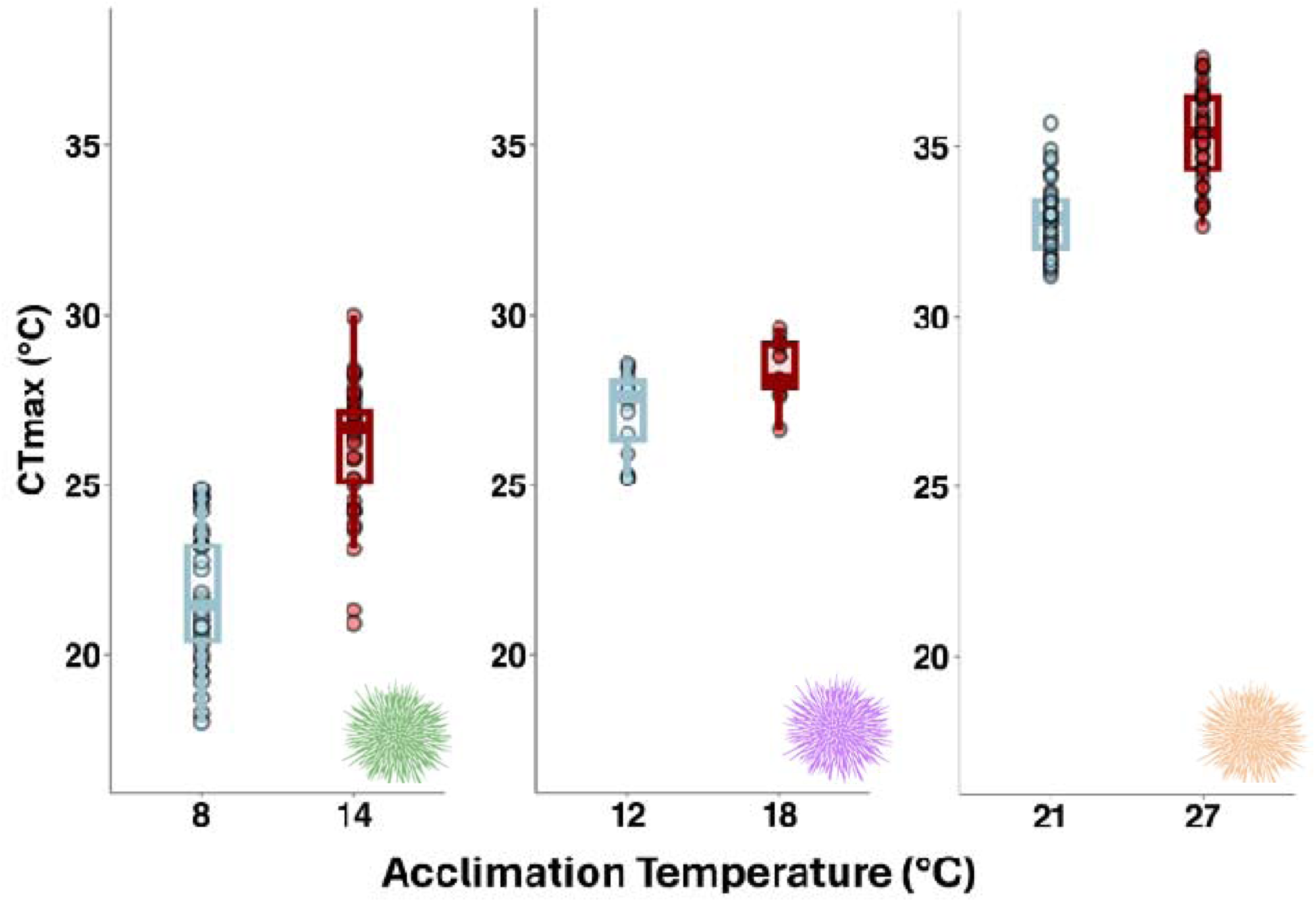
Comparisons of CT_max_ measured based on loss of adhesion in three different sea urchin species (*S. droebachiensis, S. purpuratus*, and *L. variegatus*) across two acclimation temperatures; ambient and elevated (+6°C from ambient of the species).

**Figure 3:**
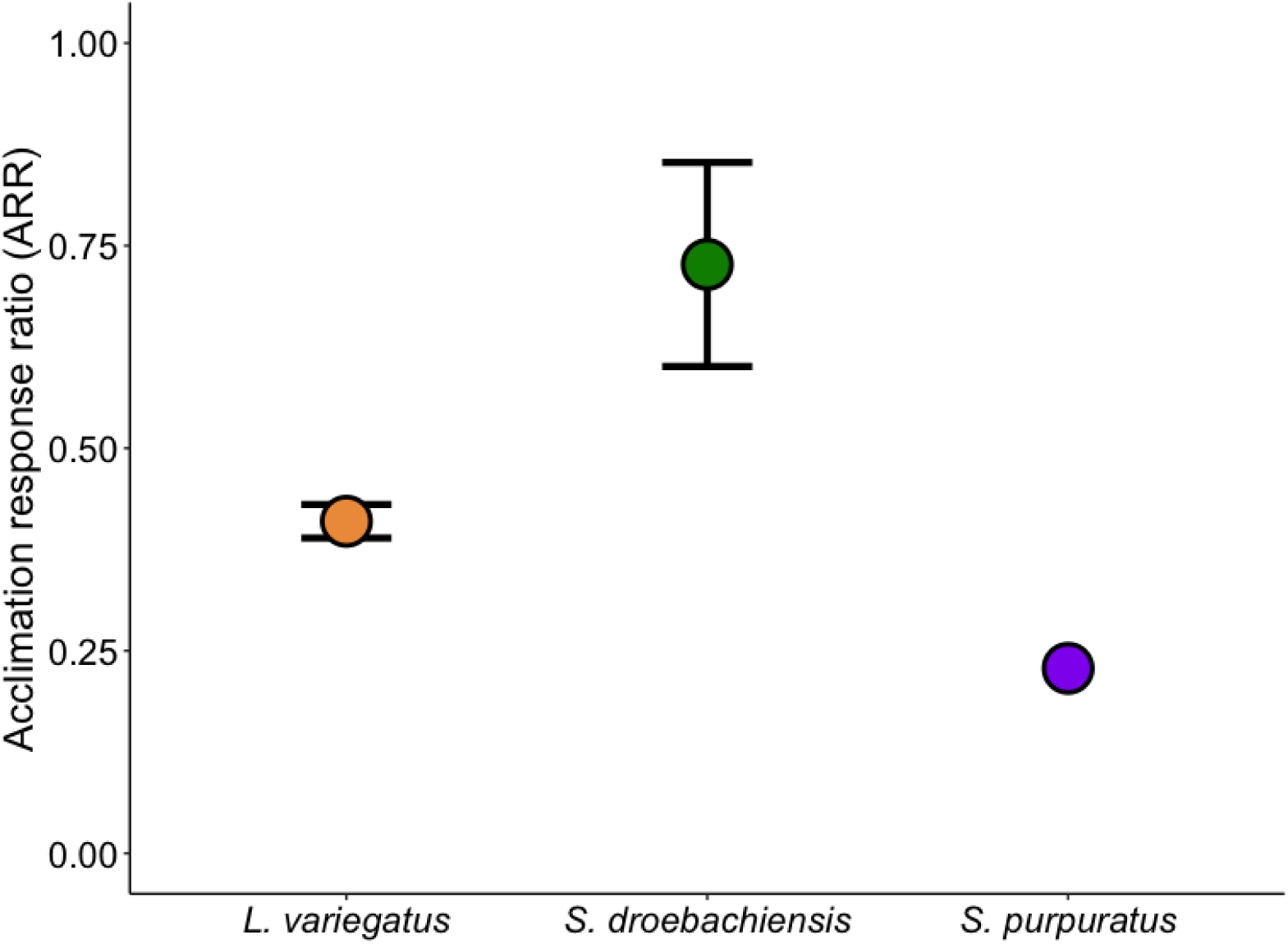
Comparison of acclimation response ratio (ARR) in three sea urchin species (*S. droebachiensis, S. purpuratus*, and *L. variegatus*) across two acclimation temperatures; ambient and elevated (+6°C from ambient of the species). Dots indicate mean and SD. No SD is plotted for *S. purpuratus* as they did not have a repeat trial.

### 3.2. Assessing repeatability and precision of the CT_max_ assay

We assessed the repeatability of our CT_max_ assay both across and within species. There was no difference between trials when repeating the CT_max_ assay for both *L. variegatus* (*F*_1,38.5_ = 0.6405, *P-value*=0.429, **Table S2; Figure 4**) and *S. droebachiensis* (*F*_1,37.6_= 1.906, *P-value*=0.176, **Table S2; Figure 4**; note: there was only one trial for *S. purpuratus*). We also found no difference between trails in spine compression or tube feet retraction for *S. droebachiensis* (**Table S2**). In contrast, *L. variegatus* differed across trials in both metrics (**Table S1**). This was likely because few *L. variegatus* individuals expressed these phenotypes and tube feet retraction was inconsistent, with frequent retraction and extension. To directly assess repeatability, we calculated repeatability scores, *R*, and found that adhesion loss had moderately high repeatability estimates within the expected values of behavioural traits: 0.466 (*D*_1,39_ = 7.47, *P-value* <0.01) for *S. droebachiensis* and 0.337 (*D*_1,77_ = 5.29, *P-value* <0.01) for *L. variegatus*. In contrast, tube feet had lower repeatability at 0.298 (*D*_1,70_=2.57, *P-value*<0.05) for *S. droebachiensis,* whilst repeatability was non-significant in *L. variegatus* (*R=*0.07, *D*_1,37_ = 0.05, *P-value* = 0.407). Lastly, spine compression had the lowest repeatability with *R* = 0 (*D*_1,31_=0, *P-value* = 1) for *S. droebachiensis*, though was surprisingly repeatable for *L. variegatus* at 0.575 (*D*_1,22_=1.36, *P-value* = 0.122), but non-significant, driven by lack of power, as only 27.5% of the urchins expressed this phenotype in both replicate trials. Additionally, we tested for correlations between the two trials to observe individual trajectories in CT_max_ (**Figure 4**) and found that correlation was higher in elevated acclimation temperature for both *S. droebachiensis* (0.539) and *L. variegatus* (0.612) compared to control treatments, 0.337 and 0.227 respectively.

**Figure 4:**
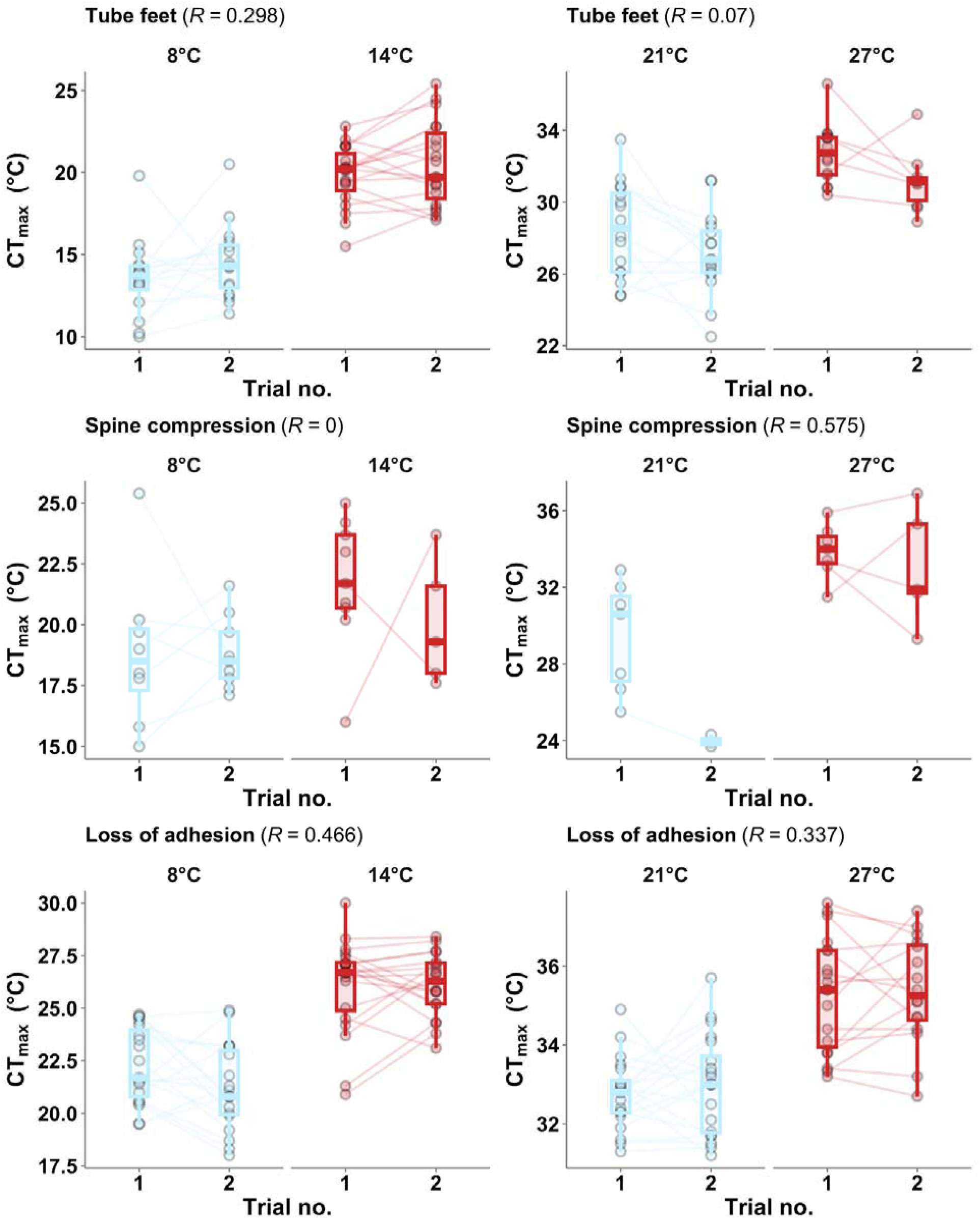
Repeatability of CT_max_ as three metrics: tube feet retraction, spine compression, and loss of adhesion for the green (*Strongylocentrotus droebachiensis*; left panels) and variegated sea urchin (*Lytechinus variegatus*; right panels) at two acclimation temperatures, ambient and elevated (+6°C) between two trials taken a week apart. Repeatability (*R*) indicated in brackets. Dots indicate individual measurements connected by individual ID between trials.

In addition, our new measure of CT_max_ by loss of adhesion had less variance than tube feet retraction and spine compression (**Table 1**) with an average of 6.7% variance compared to 13.7% and 11.1% variance, respectively. Additionally, tube feet retraction occurred less in *S. droebachiensis* (77.5%) and *L. variegatus* (87.5%). Further, spine compression was less common and did not occur in all individuals within a species or across all species. Spine compression occurred 38.75% of time for *S. droebachiensis*, 95% of the time for *S. purpuratus*, and only 27.5% of the time in *L. variegatus*. We observed on many occasions sea urchins re-erecting their spines or protracting their tube feet as temperatures continued to increase within an assay.

**Table 1:** Coefficient of variance for parameters measured during CT_max_ assay.

| Species | Loss of Adhesion | Spine compression | Tube feet |
| --- | --- | --- | --- |
| *Green* | 0.0954 | 0.1374 | 0.2101 |
| *Purple* | 0.0507 | 0.0605 | 0.0921 |
| *Variegated* | 0.0563 | 0.1362 | 0.1091 |

### 3.3. Effects of temperature acclimation on righting response

To assess performance through the acclimation and recovery periods before and after the CT_max_ assay, we used the common whole organism performance assay of righting time. To include urchins that failed to right themselves, we transformed the righting response to activity coefficient (AC). We observed no difference in temperature treatments across the weeks for both *S. droebachiensis* and *L. variegatus* (**Figure 5; Table S3**). However, for *L. variegatus*, there was a significant decline in AC across weeks, i.e., slower righting time, likely due to handling stress and starvation as the experiment progressed, particularly at the warmer ambient and elevated temperatures for this tropical species (**Figure 5; Table S3**).

**Figure 5:**
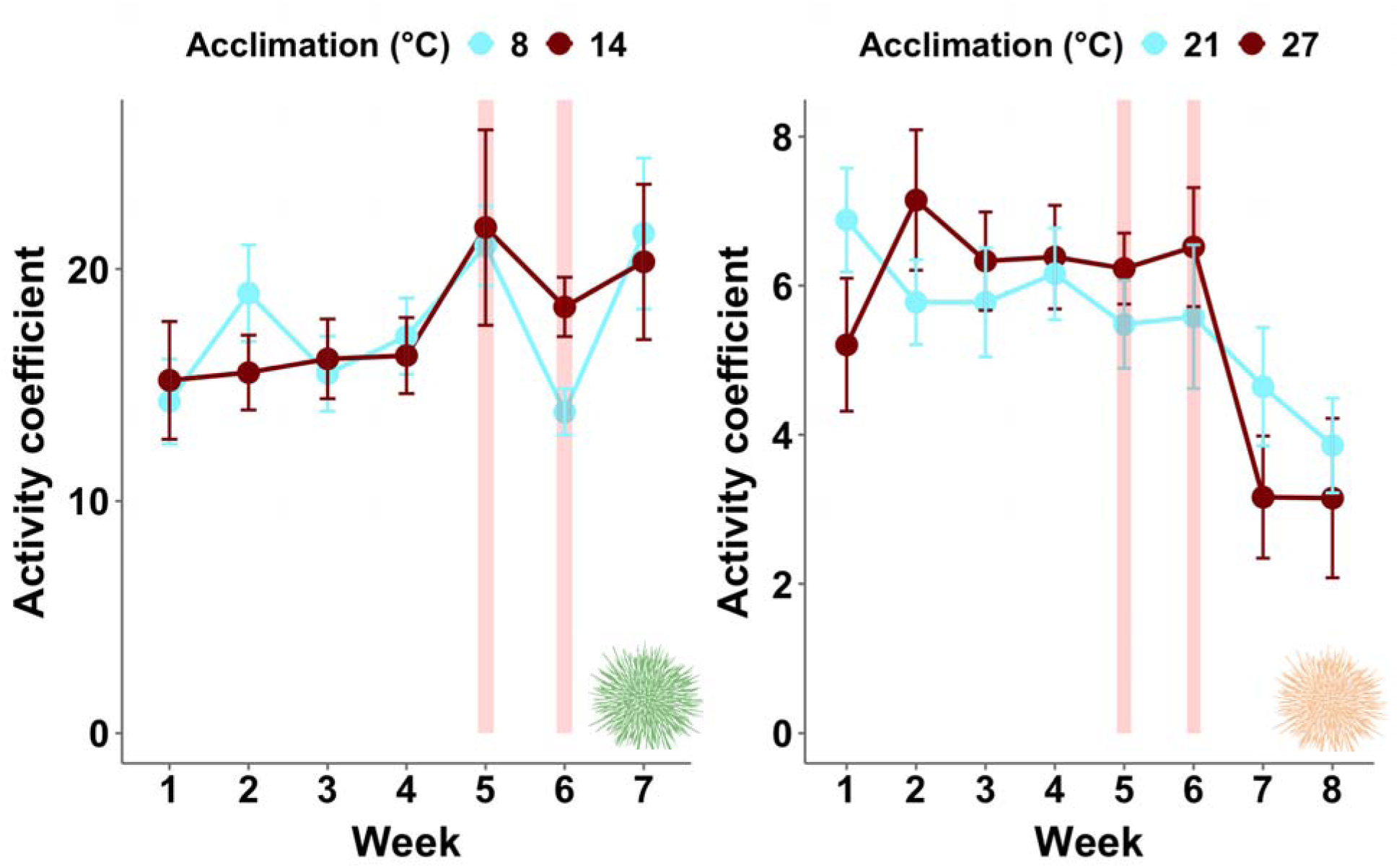
Weekly differences in activity coefficient for the green urchin (*Strongylocentrotus droebachiensis*) and the variegated urchin (*Lytechinus variegatus*) between two acclimation temperatures, ambient and elevated (+6°C). CT_max_ trial periods highlighted by red lines. Dots indicate means. Bars indicate +/- SE.

## 4. Discussion

Thermal tolerance is a key performance metric for understanding a species capacity to cope with thermal stress (Calosi et al., 2008; Pörtner et al., 2006). However, measures of thermal tolerance (e.g., CT_max_) often lack standardisation across taxa (Raby et al., 2025), which is particularly true for echinoderms, despite their ecological importance. Here, we examine the acclimation capacity of three sea urchin species from ecologically diverse habitats, develop a new adhesion-based CT_max_ assay, and compare thermal tolerance across three performance metrics. We found that acclimation to elevated temperatures (+6°C) increased thermal tolerance for all three species. Acclimation capacity was greater for both the tropical *L. variegatus* and subarctic *S. droebachiensus* relative to the latitudinally broad *S. purpuratus*, though, thermal safety margin (TSM) differed across species and treatment. We show that the new adhesion-based CT_max_ protocol is more precise and reproducible than tube feet retraction and spine compression. Our results demonstrate consistent results across a broad range of ecologically distinct taxa and contribute to our growing understanding of thermal tolerance in key foundational species.

### Acclimatory capacity differs across ecologically distinct species

Though thermal acclimation increased the CT_max_ of all three sea urchin species, consistent with the expectation that exposure to moderate elevated temperatures enhances upper thermal tolerance (Vinagre et al., 2016; Jørgenson et al., 2021; Ern et al., 2023), thermal safety margin (TSM) and acclimation response ratio (ARR) differed by species. Thermal safety margin reflects the proximity of an organism to its thermal ceiling with a smaller TSM suggesting potential greater thermal vulnerability. Under ambient conditions, *S. purpuratus* had the highest thermal safety margin (16°C), followed by the subarctic *S. droebachiensis* (13.7°C) then the tropical *L. variegatus* (11.9°C). This pattern is consistent with the climate variability hypothesis (Janzen, 1967; Stevens, 1989), which predicts that species experiencing greater variation in environmental conditions evolve broader physiological thermal tolerance. *S. purpuratus* has a broad latitudinal distribution spanning the California Current system, from Alaska to Baja California, where it experiences substantial environmental heterogeneity (Chan et al., 2017); such heterogeneity has been shown to support the evolution of a broader baseline thermal tolerance (Oliver and Palumbi 2011; Sunday et al., 2011; Rasmussen et al., 2020; Brown et al., 2024). High gene flow with pelagically dispersing larvae along this distribution may also contribute to maintaining alleles associated with thermal tolerance across populations (Pespeni et al., 2013; Pespeni and Palumbi, 2013).

In contrast, acclimation response ratio (ARR), the difference in an organism’s thermal tolerance between acclimation temperatures over the difference in acclimation temperatures (Claussen, 1977), which reflects acclimatory capacity, showed a different pattern than TSM across species. *S. droebachiensis* and *L. variegatus* had higher ARR than *S. purpuratus*. That ARR was higher in polar and tropical species than temperate species suggests there is no latitudinal pattern in acclimatory pattern. Thus, ARR, acclimatory capacity, and TSM, thermal vulnerability, capture distinct aspects of thermal physiology and are therefore not necessarily expected to covary. Consistent with this, the contrasting responses among species suggest that high baseline thermal tolerance and strong acclimatory capacity are not necessarily coupled.

The relatively small increase in CT_max_ following warm acclimation and correspondingly low ARR of S. purpuratus may indicate limited scope for further plastic increases because this species already possesses a comparatively high thermal ceiling under ambient conditions. In contrast, both *S. droebachiensis* and *L. variegatus* showed high capacity to raise their upper thermal capacity, and indeed, *S. droebachiensis* had an ARR of 0.73, nearly three times the reported average for marine ectotherms (Gunderson and Stillman, 2015). These results are consistent with Richard et al. (2012), who found that S. droebachiensis acclimated at 10.3°C had greater upper thermal tolerance than individuals acclimated at 7.1°C, and demonstrates that this response persists across a larger acclimation-temperature difference approaching the upper range of marine heatwave conditions (Oliver et al., 2018; Sen Gupta et al., 2020). More broadly, the absence of a consistent relationship between acclimation capacity and latitude agrees with Gunderson and Stillman (2015) and with studies reporting limited latitudinal variation in CT_max_ among aquatic ectotherms (Comte and Olden, 2017; Sunday et al., 2012). It is important to note that these results are representative of only one population within each species, and different populations may have different acclimatory across latitudinal clines (Castañeda et al., 2015; Healy et al., 2019; Sasaki et al., 2024). Moreover, in the present study we use adult urchins, whilst larvae and juveniles may be more thermally sensitive and can experience different selective pressures across ontogeny (Collin et al., 2021; Nunez et al., 2026). Integrating thermal limits across eggs, larvae, juveniles, and adults will therefore be essential for predicting how species-level acclimation capacity translates into population persistence under future global change.

### Creating an assay to measure thermal tolerance

A crucial aim of this study was to generate a replicable, precise proxy of CT_max_. We achieved this by assessing loss of adhesion within a flowthrough cage, which we showed to be repeatable and had less variability than other proxies measured. A method used by some urchin studies was developed by Hernandez et al., (2004), where they measure phenotypic responses in stages including movement towards the bottom of the aquaria, retraction of the tube feet and relaxation of the spines, where they demonstrated significant differences in the CT_max_ measured during a five-stage process in the red sea urchin (*Mesocentrotus franciscanus*). However, we found in the three species of urchins in the present study that spine compression and tube feet were often inconsistent and unreliable, with individuals sometimes not presenting either phenotype at all during the assay or expressing the phenotype multiple times at different stages with high individual variation. Loss of adhesion is also biologically relevant as it is indicative of loss of motor function and reduced predator escape potential (Lutterschmidt and Hutchison, 1997). For urchins, and many other intertidal and subtidal species, adhesion also represents capacity to remain attached to substrate during strong wave action or in strong currents and the ability to resist predation by a fish or otter, for example (Dayton 1973,1975; Denny 1988; Santos et al., 2009). We also found that righting response, a proxy for whole organism performance commonly used in urchins (Brothers and McClintock, 2015; Percy, 1973) that indicates neuromuscular activity (Lawrence and Cowell, 1996), did not differ between acclimation temperatures or through time across the duration of the study. These results suggest that the CT_max_ assay itself did not impact organismal performance, as is supported by the repeatability of the assay with little mortality. As CT_max_ is strongly dependent on methodological context (Terblanche et al., 2007), we propose this method as a standard measure of CT_max_ in sea urchins going forward.

### Future directions in understanding thermal tolerance across marine taxa

Though this assay provides a strong methodology to assess thermal tolerance in sea urchins, additional studies on other taxa and populations would broaden the comparative framework of this thermal tolerance assay. Beyond our present assay on sea urchins, loss of attachment has already been used to determine upper thermal tolerances in molluscs (Salas et al., 2014; Yu et al., 2021), but our design could be expanded to other adhering taxa, assessing the impacts of acclimation duration and heat stress carryover effects, and testing multiple populations and life stages. Additionally, it would be valuable to determine how persistent carryover effects are and whether they translate into longer-term reductions in organismal fitness; where here we were limited by a five week assay (4-week acclimation period prior to CT_max_ and righting response after one week of recovery). Survival is a key measure of organismal fitness; Richard et al. (2012) reported increased mortality in Arctic sea urchins, molluscs, and amphipods following thermal stress, demonstrating long-term effects of acute thermal stress. Here we show that *L. variegatus* had strong TSM and ARR, however, other studies have shown weaker responses in an exclusively tropical species of urchin (Collin et al., 2018), so *L. variegatus* may be more thermally tolerant due to its wider range of habitat from Brazil up to the sub-tropics in North Carolina (Moore et al., 1963). It is also pertinent to note that thermal tolerance is lower at the larval stage of many marine ectotherms (Collin et al., 2021). It is also important to consider the transgenerational impact of thermal stress where heatwave events experienced by parents could drive acclimation to thermal stress in larval offspring. Beyond upper thermal tolerance, lower thermal tolerance (CT_min_) may be much more variable among species (Collin et al., 2018; Sunday et al., 2012) and relevant as cold snaps become more frequent and upwelling events drive an increase in cold water coming up from depth to the sub- and intertidal zones. However, CT_min_ was not possible to quantify in the present study because in pilot experiments both green and purple sea urchins were still moving and adhering at our lowest experimental temperatures, sub-zero (-2°C). Together, these limitations highlight that short-term increases in adult CT_max_ represent one component of climate resilience. Resolving how thermal tolerance varies among species, across populations and life stages, and through time will be critical for determining whether acclimation can buffer populations against future global change.

### Conclusions

We show that sea urchins from ecologically diverse habitats all acclimate to elevated thermal stress but differ in their acclimation capacity. *S. purpuratus* exhibited the highest baseline thermal safety margin yet had the weakest acclimatory response. In contrast, the subarctic *S. droebachiensis* and tropical *L. variegatus* displayed a greater capacity to raise CT_max_ when exposed to elevated temperatures, highlighting the need to understand thermal tolerance across ecologically diverse taxa. We showed our new CT_max_ assay to be repeatable and validated results across species. However, it is important to further test the assay with other taxa to understand the diversity of patterns observed in thermal tolerance. Our results contribute to the growing understanding of thermal tolerance in marine ectotherms and could be used to inform predictions on species performance under rapidly occurring global change. This study provides a new, precise methodology that can help expand our understanding of the survival capacity for a taxonomic group that is foundational in many ecosystems.

## Acknowledgements

This work was supported by the National Science Foundation award IOS 1943396 to M.H.P. This work was also supported by start-up funds from the University of Vermont to J.C.B.N. D.E.S was supported by the Office of the Vice President for Research at the University of Vermont. The authors thank the Vermont Advanced Computing Center (VACC; https://www.uvm.edu/vacc) for providing computational resources that contributed to this publication.

## Data availability

All data and code available at https://github.com/DanielSadler96/Sea_Urchin_CTmax

