## Supplemental Information for "A novel CT_max_ assay reveals divergent thermal acclimation capacity across three ecologically distinct sea urchins"

**Contents:**

S2	Table S1

S3	Table S2

S4	Table S3

**Table S1:** List of components to create the CTmax assay system

| **Section** | **Materials** |
| --- | --- |
| ***Cage*** | Perspex sheet (3mm thick) laser cut into 5 cm x 5 cm |
|  | Alegi Aquaria grating (10 mm x 10 mm grates) cut into 5 cm x 5 cm squares |
|  | Zipties (black for structure, coloured for identification) |
| ***CTmax system*** | 30 L open-top plastic storage container (51.75 cm x 35.88 cm x 15.24 cm) |
|  | Natural cotton insulating material (Frost King) |
|  | Inkbird WiFi ITC-308 |
|  | 500 W Hygger 802 Aquarium Titanium Heater |
|  | Testo 112 NTC Food Thermometer with Waterproof immersion probe NTC |
|  | 5 W water pump |
|  | 1 inch airstone attached to Tetra whisper air pump |

**Table S2:** Statistical summaries of CT_max_ measures across all species of sea urchin measured. Models were linear mixed effect models (LMM) with the exception of the purple sea urchin (*Strongylocentrotus purpuratus*) which was a standard linear model due to the absence of repeated measures.

| ***Species*** | ***Trait*** | ***Variable*** | ***F*** ***Value*** | ***P*** ***value*** |
| --- | --- | --- | --- | --- |
| **Green**  *+(1\|Urchin ID)* | Adhesion | Treatment | 70.9522 | **5.227x10^-10^***** |
|  |  | Trial | 1.906 | 0.1758 |
|  | Spines | Treatment | 6.1723 | **0.0192*** |
|  |  | Trial | 0.6415 | 0.4299 |
|  | Tube feet | Treatment | 101.5357 | **2.787x10^-12^***** |
|  |  | Trial | 3.6522 | 0.0640 |
| **Purple**  (no repeated trials) | Adhesion | Treatment | 4.2547 | **0.04603*** |
|  | Spines | Treatment | 23.646 | **3.432x10^-5^***** |
|  | Tube feet | Treatment | 22.946 | **2.551x10^-5^***** |
| **Variegated**  *+(1\|Urchin ID)* | Adhesion | Treatment | 56.1855 | **4.15x10^-9^***** |
|  |  | Trial | 0.6405 | 0.4285 |
|  | Spines | Treatment | 16.281 | **3.739x10^-5^***** |
|  |  | Trial | 6.2976 | **0.02572*** |
|  | Tube feet | Treatment | 45.3673 | **5.47x10^-8^ ***** |
|  |  | Trial | 8.4079 | **0.006586**** |

**Table S3:** Summary of statistical analysis for righting response across green and variegated urchins using glmmTMB models, presented as analysis of deviance (Type II Chisq)

| ***Species*** | ***Variable*** | ***Chisq*** | ***Df*** | ***P value*** |
| --- | --- | --- | --- | --- |
| **Green** | Temperature | 0.0118 | 1 | 0.9133 |
|  | Week | 18.4494 | 6 | **0.0052**** |
| **Variegated** | Temperature | 0.0006 | 1 | 0.9802 |
|  | Week | 26.0247 | 7 | 0.0005*** |
